# Genome-scale enzymatic reaction prediction by variational graph autoencoders

**DOI:** 10.1101/2023.03.08.531729

**Authors:** Cheng Wang, Chuang Yuan, Yahui Wang, Ranran Chen, Yuying Shi, Gary J. Patti, Qingzhen Hou

## Abstract

**Background:** Enzymatic reaction networks are crucial to explore the mechanistic function of metabolites and proteins in biological systems and understanding the etiology of diseases and potential target for drug discovery. The increasing number of metabolic reactions allows the development of deep learning-based methods to discover new enzymatic reactions, which will expand the landscape of existing enzymatic reaction networks to investigate the disrupted metabolisms in diseases.

**Results:** In this study, we propose the MPI-VGAE framework to predict metabolite-protein interactions (MPI) in a genome-scale heterogeneous enzymatic reaction network across ten organisms with thousands of enzymatic reactions. We improved the Variational Graph Autoencoders (VGAE) model to incorporate both molecular features of metabolites and proteins as well as neighboring features to achieve the best predictive performance of MPI. The MPI-VGAE framework showed robust performance in the reconstruction of hundreds of metabolic pathways and five functional enzymatic reaction networks. The MPI-VGAE framework was also applied to a homogenous metabolic reaction network and achieved as high performance as other state-of-art methods. Furthermore, the MPI-VGAE framework could be implemented to reconstruct the disease-specific MPI network based on hundreds of disrupted metabolites and proteins in Alzheimer’s disease and colorectal cancer, respectively. A substantial number of new potential enzymatic reactions were predicted and validated by molecular docking. These results highlight the potential of the MPI-VGAE framework for the discovery of novel disease-related enzymatic reactions and drug targets in real-world applications.

**Data availability and implementation:** The MPI-VGAE framework and datasets are publicly accessible on GitHub https://github.com/mmetalab/mpi-vgae.

**Author Biographies:** **Cheng Wang** received his Ph.D. in Chemistry from The Ohio State Univesity, USA. He is currently a Assistant Professor in School of Public Health at Shandong University, China. His research interests include bioinformatics, machine learning-based approach with applications to biomedical networks.

**Chuang Yuan** is a research assistant at Shandong University. He obtained the MS degree in Biology at the University of Science and Technology of China. His research interests include biochemistry & molecular biology, cell biology, biomedicine, bioinformatics, and computational biology.

**Yahui Wang** is a PhD student in Department of Chemistry at Washington University in St. Louis. Her research interests include biochemistry, mass spectrometry-based metabolomics, and cancer metabolism.

**Ranran Chen** is a master graduate student in School of Public Health at University of Shandong, China.

**Yuying Shi** is a master graduate student in School of Public Health at University of Shandong, China.

**Gary J. Patti** is the Michael and Tana Powell Professor at Washington University in St. Louis, where he holds appointments in the Department of Chemisrty and the Department of Medicine. He is also the Senior Director of the Center for Metabolomics and Isotope Tracing at Washington University. His research interests include metabolomics, bioinformatics, high-throughput mass spectrometry, environmental health, cancer, and aging.

**Leyi Wei** received his Ph.D. in Computer Science from Xiamen University, China. He is currently a Professor in School of Software at Shandong University, China. His research interests include machine learning and its applications to bioinformatics.

**Qingzhen Hou** received his Ph.D. in the Centre for Integrative Bioinformatics VU (IBIVU) from Vrije Universiteit Amsterdam, the Netherlands. Since 2020, He has serveved as the head of Bioinformatics Center in National Institute of Health Data Science of China and Assistant Professor in School of Public Health, Shandong University, China. His areas of research are bioinformatics and computational biophysics.

**Key points:**

- Genome-scale heterogeneous networks of metabolite-protein interaction (MPI) based on thousands of enzymatic reactions across ten organisms were constructed semi-automatically.
- An enzymatic reaction prediction method called Metabolite-Protein Interaction Variational Graph Autoencoders (MPI-VGAE) was developed and optimized to achieve higher performance compared with existing machine learning methods by using both molecular features of metabolites and proteins.
- MPI-VGAE is broadly useful for applications involving the reconstruction of metabolic pathways, functional enzymatic reaction networks, and homogenous networks (e.g., metabolic reaction networks).
- By implementing MPI-VGAE to Alzheimer’s disease and colorectal cancer, we obtained several novel disease-related protein-metabolite reactions with biological meanings. Moreover, we further investigated the reasonable binding details of protein-metabolite interactions using molecular docking approaches which provided useful information for disease mechanism and drug design.

## 1. Introduction

Characterizing enzymatic reactions is important to understand biochemical transformations, allosteric inhibition, and protein signaling [1–3]. Enzymatic reactions start with interactions between metabolites and proteins that occur at the active site of enzymes and are building blocks for metabolic networks [4, 5]. Functional annotations and characterizations of enzymatic reactions pave the way to understanding the metabolic mechanisms and uncovering associations between metabolomics and diseases in biomedical research [6, 7]. The rapid advances in high-throughput metabolomics and proteomics technologies promote the systematic profiling of metabolites and proteins [8–10]. The discovery of novel enzymatic reactions is essential to investigating their mechanistic roles in cellular metabolisms and disease progression. Experimental approaches have been developed to systematically discover enzymatic reactions by mapping metabolite-protein interactions (MPI) in cells [11–13]. For example, 1678 interaction pairs between 20 designated metabolites and cellular proteins in *Escherichia coli* were experimentally determined by mass spectrometry (MS) [11]. High-resolution NMR relaxometry was recently developed to detect MPIs in biological fluids [14]. Though these experimental methods offer high reproducibility and accuracy to characterize enzymatic reactions, the low binding affinity of MPIs and labor-intensive sample preparation hamper the process for large-scale enzymatic reaction screening.

Since the formation of an enzyme-substrate complex is a prerequisite for an enzymatic reaction, accurate prediction of metabolite-protein interactions would facilitate the discovery of new enzymatic reactions. Recently, a variety of machine learning-based methods have been developed to predict MPI computationally, while most of these methods focus on predicting the allosteric interaction instead of the likelihood of enzymatic reaction [15, 16]. It is imperative to develop efficient computational approach to predict MPI for enzymatic reaction. Thousands of enzymatic reactions have been cataloged in multiple metabolic databases such as KEGG, Reactome, and PathBank [17–19]. These interconnected graph-based representations of enzymatic reactions generate genome-scale metabolic networks, while there are still no computational methods to fully explore the metabolic reaction networks for MPI prediction.

The prediction of MPI based on the enzymatic reaction network takes advantage of inter-connected features of proteins and metabolites, which can be formulated as a so-called “link prediction” computational problem [20, 21]. Graph neural networks (GNNs) integrate the graph topology and node/edge features and show superior performance of link prediction than traditional machine learning methods [22–24]. Graph neural networks have been widely applied to recognize the link in network properties of biomolecules ranging from protein structure prediction, protein-protein interaction networks, and protein-RNA binding, to multi-omics disease studies [25–30]. However, there are no available GNN methods for MPI prediction within enzymatic reaction network.

In this study, we constructed a heterogeneous network of metabolite-protein functional interaction networks from thousands of enzymatic reactions and developed a Variational Graph Autoencoders (MPI-VGAE) framework to predict enzymatic reactions from different organisms. Ten organism-wise MPI networks were constructed and MPI prediction with the VGAE model was trained and optimized. By comparing with conventional similarity-based and graph-based methods, we demonstrated that the MPI-VGAE method out- performed other models for MPI prediction in different genome-scale MPI networks with the highest AUC and AP scores. The heterogeneous node features of metabolites and proteins and neighboring information were well incorporated into the MPI-VGAE framework via the feature transformation module. We applied MPI-VGAE framework to multiple scenarios, including reconstruction of metabolic pathways, functional metabolic networks and homogeneous metabolic reaction networks. Finally, the MPI-VGAE framework was applied to study Alzheimer’s disease and colorectal cancer to reconstruct the MPI network by hundreds of disrupted metabolites and proteins. MPI-VGAE could predict new potential enzymatic reactions, which were further investigated with the possible binding poses for several examples. The MPI-VGAE framework will facilitate the discovery of novel enzymatic reactions in biomedical research.

## 2. Methods

### 2.1 Datasets and characteristics of proteins and metabolites

To construct a metabolite-protein interaction (MPI) network, the metabolite-protein interactions were extracted from all metabolic pathways in PathBank. Each metabolic pathway contains metabolites and protein information. The metabolite-protein interaction networks were constructed and specified for ten organisms separately, such as *Homo sapiens*. In the MPI network, each metabolite and protein were modeled as a node and each interaction was modeled as an edge. The classes of proteins and metabolites were characterized and classified by using BRENDA and RefMet. The dataset curation process is depicted in Figure 1A. The homogenous metabolic reaction network was constructed based on the chemical reactions in the KEGG database. Metabolites in all pathways of different organisms in the KEGG database were extracted via KEGG API. In the metabolic reaction network, each metabolite was modeled as a node and each metabolic reaction was modeled as an edge.

**Figure 1.**
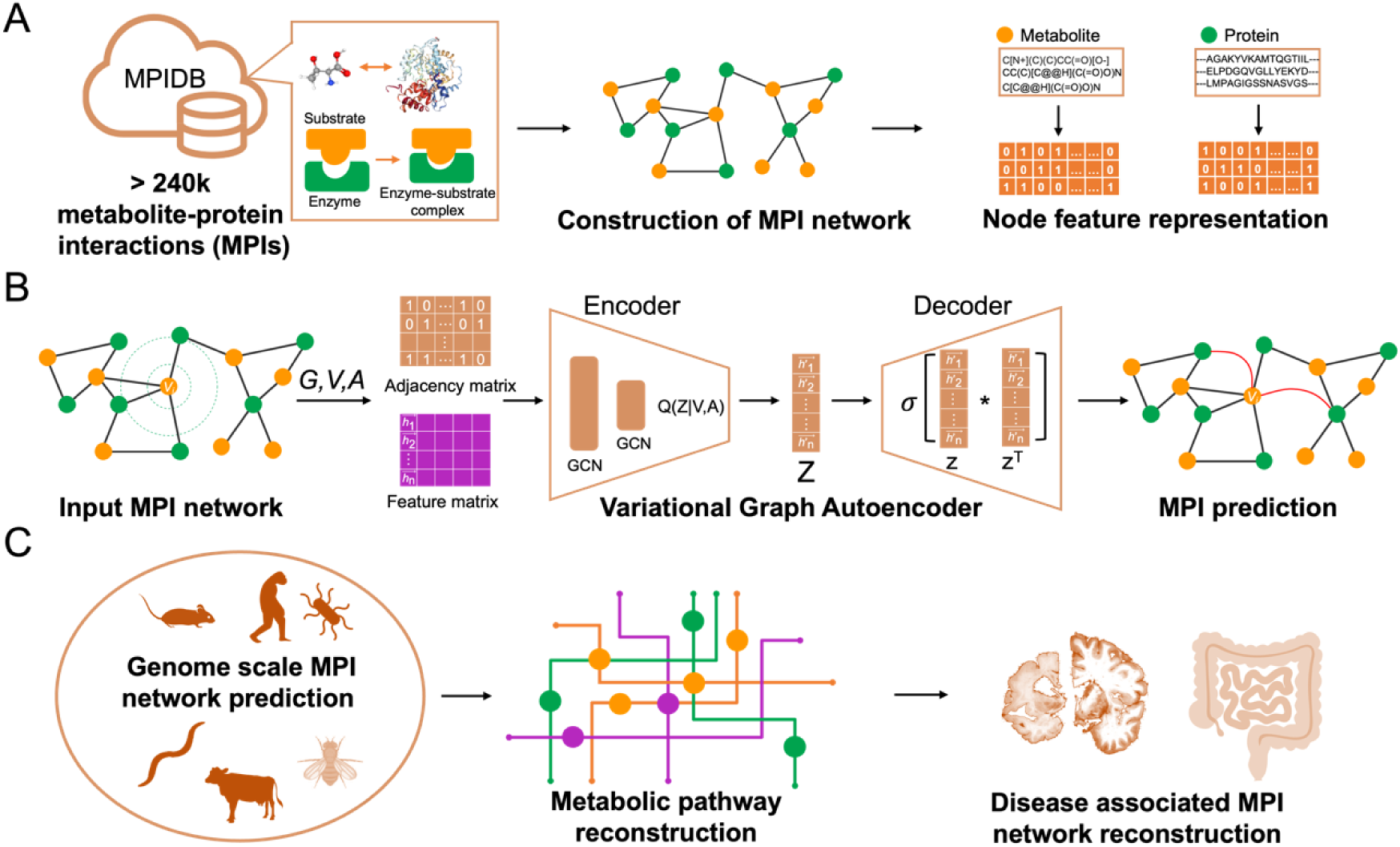
Overview of metabolite-protein interactions prediction by Variational Graph Autoen-coder. Panel A depicts the construction of the metabolite-protein interaction (MPI) network and featurization of metabolites and proteins. Panel B depicts the Variational Graph Autoencoder algorithm to predict MPIs. Adjacency matrix and feature matrix were constructed based on the MPI network and explicit features of metabolites and proteins. The encoder module takes both the adjacency matrix and feature matrix by two graph convolutional layers, followed by the decoder for the reconstruction of the adjacency matrix. The likelihood of MPI is computed based on the latent embedding vectors. Panel C depicts the application scenarios of MPI-VAGE framework, including genome scale MPI prediction, metabolic pathway reconstruction, and reconstruction of disease associated MPI networks.

### 2.2 Feature representation of metabolites and proteins

The vectorized features of metabolites and proteins were generated as input for the graph-based neural network models. For metabolites, molecular fingerprints were used to represent the features of each metabolite. A molecular fingerprint uses a series of binary digits to indicate the presence or absence of a particular substructure in the molecule. Three popular molecular fingerprint representations were considered for comparison, including the Extended Connectivity Fingerprint (ECFP), MACCS keys, and topological fingerprints [31, 32]. RDKit (https://www.rdkit.org/) was used to generate molecular fingerprints based on the SMILES string of each molecule, which yielded a binary vector with a fixed-length [33]. In addition, the fingerprints of metabolites were further normalized and transformed by using principal component analysis (PCA). Essentially, all fingerprints of 78,726 metabolites were fit by using PCA and transformed into a 1024-bit vector. For proteins, the raw amino-acid sequence was used to capture the feature information and converted into a vector with a length of 1024 based on a pre-trained SeqVec model and an ESM-1b transformer model [34, 35]. The SeqVec model is a representation learning model based on the language model ELMo, taken from natural language processing. ELMo creates embeddings in 0.03s per protein sequence, on average. The state-of-the-art ESM-1b transformer protein language model is a deep contextual language model trained on 86 billion amino acids across 250 million protein sequences spanning evolutionary diversity. The generated features were used as input for graph neural network-based model training and optimization.

### 2.3 Mathematical representation of metabolic reaction graph

The metabolite-protein interaction network and metabolic reaction network were represented as undirected graph G = (V, A, X) with N = |V| nodes. For each graph, G was represented by its adjacency matrix A ∈ R^N×N^. A ∈ {1,0}^N×N^, where 1 denotes there is an edge (enzymatic reaction) between a pair of nodes (metabolite/protein) and 0 otherwise. Given that the reaction direction was not considered in this study, the graphs were undirected and no weights between edges were specified. The adjacency matrix A was symmetric and unweighted. The feature matrix X∈R^N×D^ denotes the node features. The node features consist of both explicit and latent features. Explicit features are node attributes, such as the molecular fingerprints of metabolites and vectorized representation of proteins. Latent features are the matrix representations of the graph learned by the graph-embedding methods. These low-dimensional latent representations preserve the properties of the graph such as the local neighborhood of nodes. In the current study, the latent features were used in the graph-embedding models, including Node2vec and Variational Graph Autoencoders.

### 2.4 Variational Graph Autoencoders Model

The variational graph autoencoders model is an unsupervised learning framework for graph-structured data using variational Bayesian methods. Here, we recapitulate the VGAE model and illustrate how it incorporates the metabolite and protein features for link prediction. The VGAE model first maps the nodes onto low-dimensional vector features by an encoder and uses a decoder to reconstruct the original graph topological information. The encoder of VGAE simultaneously incorporates both node structural information and attributes by two graph convolutional layers, as shown below:

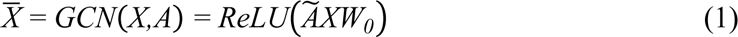

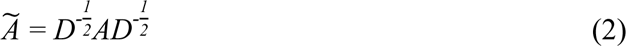

where *Ã* is the symmetrically normalized adjacency matrix. The graph convolutional network (GCN) layer performs convolution on graphs to extract local substructure features for individual nodes. Then it aggregates node-level features into a graph-level feature vector. The latent distribution is then produced by the encoder as following:

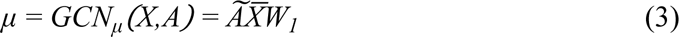

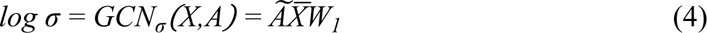

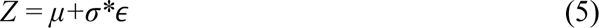

where ɛ∼𝖭(0,1) and Z denotes the graph-embedding matrix. Then, the encoder is formulated as:

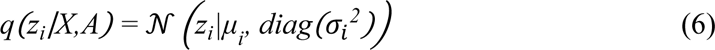

The decoder decodes the embeddings by reconstructing the graph adjacency matrix:

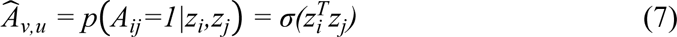

The loss function is a combination of the reconstruction loss and the KL-divergence:

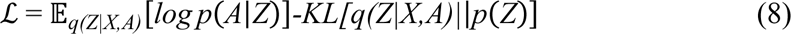

The workflow of the VGAE model for metabolite-protein interaction prediction is shown in Figure 1B. First, a graph of metabolite-protein interactions is constructed. The features of metabolites and proteins are also embedded. Second, the VGAE model learns the embedding vectors of metabolites and proteins. Lastly, the interaction of a pair of metabolites and proteins is predicted based on the learned embedding vectors.

### 2.5 Model training and performance evaluation

During the model training, all the existing edges in the network were considered as positive examples while all non-existent edges in the network were considered as negative examples. Due to the sparseness of the MPI and metabolic reaction network, the number of true positive examples was significantly less than the number of true negative examples. This would cause an imbalance problem during the training process. We applied the undersampling approach to balance the dataset by considering an equal number of positive and negative examples during the model training and testing progress. All the positive and negative edges in the original graph of each organism was split into training and testing datasets with a proportion of 80% and 20%, respectively. The training dataset was used for feature selection, model training, and optimization. 5-fold cross-validation was used for the optimal feature selection and optimization of parameters. To measure the performance of the model, the Area Under the Curve (AUC) Receiver Operating Characteristics (ROC) and Precision-Recall (PR) curve were used. The ROC curve was plotted with the true positive rate (TPR) against the false positive rate (FPR), where TPR is on the y-axis and FPR is on the x-axis. The higher the AUC, the better the performance of the model at predicting positive and negative links. The Precision-Recall (PR) curve was constructed by calculating and plotting the precision and the recall at a variety of thresholds. The evaluation functions are listed as follows:

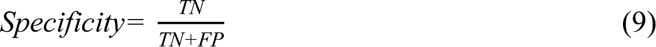

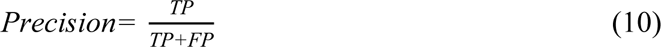

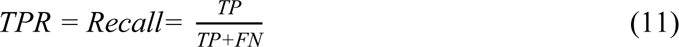

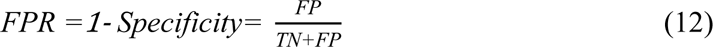

where TP stands for true positive (i.e., the model predicts a link exists between a pair of nodes and there is a reaction between a protein and a metabolite in the dataset), FP stands for false positive, TN stands for true negative (i.e., the model predicts no link exists between a pair of nodes and there is no reaction between a protein and a metabolite in the dataset), and FN stands for false negative.

### 2.6 Model comparison and evaluation

We compared the VGAE model with several baseline machine learning models, including similarity-based models, random walk-based models, and graph-embedding models. The similarity-based measure is a type of unsupervised approach that computes the likelihood of each non-existing link from the similarity score of two nodes, such as Adamic-Adar and Preferential Attachment [36, 37]. Spectral clustering was also included in the baseline models [38]. For graph data, spectral clustering creates node representations by taking top *d* eigenvectors of the normalized Laplacian matrix of the graph. Then, the feature vector representations of nodes were used for computing the likelihood of a pair of nodes. Random walk-based methods learn node representations by generating “node sequences” through random walks in graphs, as inspired by Natural Language Processing (NLP), which tries to learn word representations from sentences. Node2vec is a skip-gram-based approach to learning node embeddings from random walks within a given graph [39]. Graph embedding is a graph representation learning technique that converts graph data into vectors followed by generating a representation of nodes in a lower-dimensional space. Since the embeddings preserve the graph properties such as the local neighborhood of nodes, the likelihood of a link between two targeted nodes is computed based on the embeddings of the nodes. GraphSAGE is an inductive graph neural network model that incorporates node feature information to efficiently generate representations on large graph, which was used to as benchmark model to compare MPI-VGAE [40].

### 2.7 Molecular docking for metabolite-protein interactions

AutoDock Vina was implemented for protein-metabolites docking. We selected the predicted enzyme reactions of protein Cholesterol side-chain cleavage enzyme (CYP11A) and Aldo-keto reductase family 1 member C4 (AKR1C4) binding with 24-Hydroxycholesterol and 27-Hydroxycholesterol respectively as examples. For CYP11A, we used the protein structure from Protein Data Bank (PDB ID 3N9Z) as starting structure. The original binding ligands (22-hydroxycholesterol and Adrenodoxin) of 3N9Z were depleted and the interacting sites between ligands and protein were calculated by distance measure (distance <6 angstrom). We then built the simulation box covering all interaction regions and perform protein-metabolites docking between CYP11A with 24-Hydroxycholesterol or 27-Hydroxycholesterol respectively using AutoDock Vina. The docking processes were run for ten times with random seeds and the conformation with lowest binding affinity was selected as the representing structure.

For protein AKR1C4, we downloaded the predicted structure from Alphafold2 Database (averaged pLDDT > 90) and constructed simulation box covering all proteins to find the most possible binding pose. The docking was also performed between AKR1C4 structure and 24-Hydroxycholesterol or 27-Hydroxycholesterol respectively for 10 times. The structure with lowest binding affinity was selected.

## 3. Results

### 3.1 Overview of the MPI-VGAE framework

The complete MPI-VGAE framework is depicted in Figure 1. The genome-scale metabolite-protein interaction networks for different organisms were automatically curated based on thousands of metabolic pathways from PathBank, followed by manual screening to remove redundant nodes and edges. Explicit node features included molecular fingerprints of metabolites and sequence-based features of proteins, which formed a feature matrix. The Variational Graph Autoencoder model was trained and optimized by using the adjacency matrix and feature matrix from the MPI network. The encoder module consisted of double graph convolutional layers, followed by the decoder for the reconstruction of the adjacency matrix. The likelihood of MPI was computed based on the embedding vectors.

### 3.2 Characteristics of metabolite-protein interaction networks

The details of metabolites and proteins in the MPI network are shown in Figure 2. For metabolites, fatty acyls (FA), organic acid (OA), and nucleic acid (NA) are the most prevalent metabolite classes, which account for 46% of the total (Figure 2A). Among the seven enzyme classes (i.e., oxidoreductase, ligase, hydrolase, isomerase, lyase, transferase, and translocase), transferase and oxidoreductase are the two main enzyme categories in all metabolic pathways (Figure 2B), which suggest that most metabolic reactions involve group transfer reactions and oxidation-reduction reactions. The node degree represents the number of interconnected nodes in the MPI network. Based on the node degree distributions in the integrated MPI network of all organisms in Figure 2C and D, the median links of metabolites and proteins are 3 and 5, respectively. Among all metabolites, 55.4% have degrees more than three, suggesting that more than half of the metabolites participate in at least three enzymatic reactions. Among all proteins, 78.5% have degrees more than three, suggesting that most proteins may catalyze reactions that involve three or more unique metabolite substrates. There are 258 metabolites (11.2%) and 256 proteins (10.5%) that have degrees more than ten. It should be noted that the node degree distributions may vary across different organisms because of the variation of metabolic pathways among different organisms. The details of the metabolite-protein interaction network for each organism are summarized in Table 1. The MPI network was visualized by using the Fruchterman-Reingold layout. Figure 3 shows an example of the human MPI network. The top interconnected metabolites and proteins with the highest degree are annotated. Essential metabolites such as ATP, NAD, and L-glutamic acid are the most interconnected metabolites in the MPI network. Cytochromes P450 family enzymes have the most diverse connections with metabolites.

**Figure 2.**
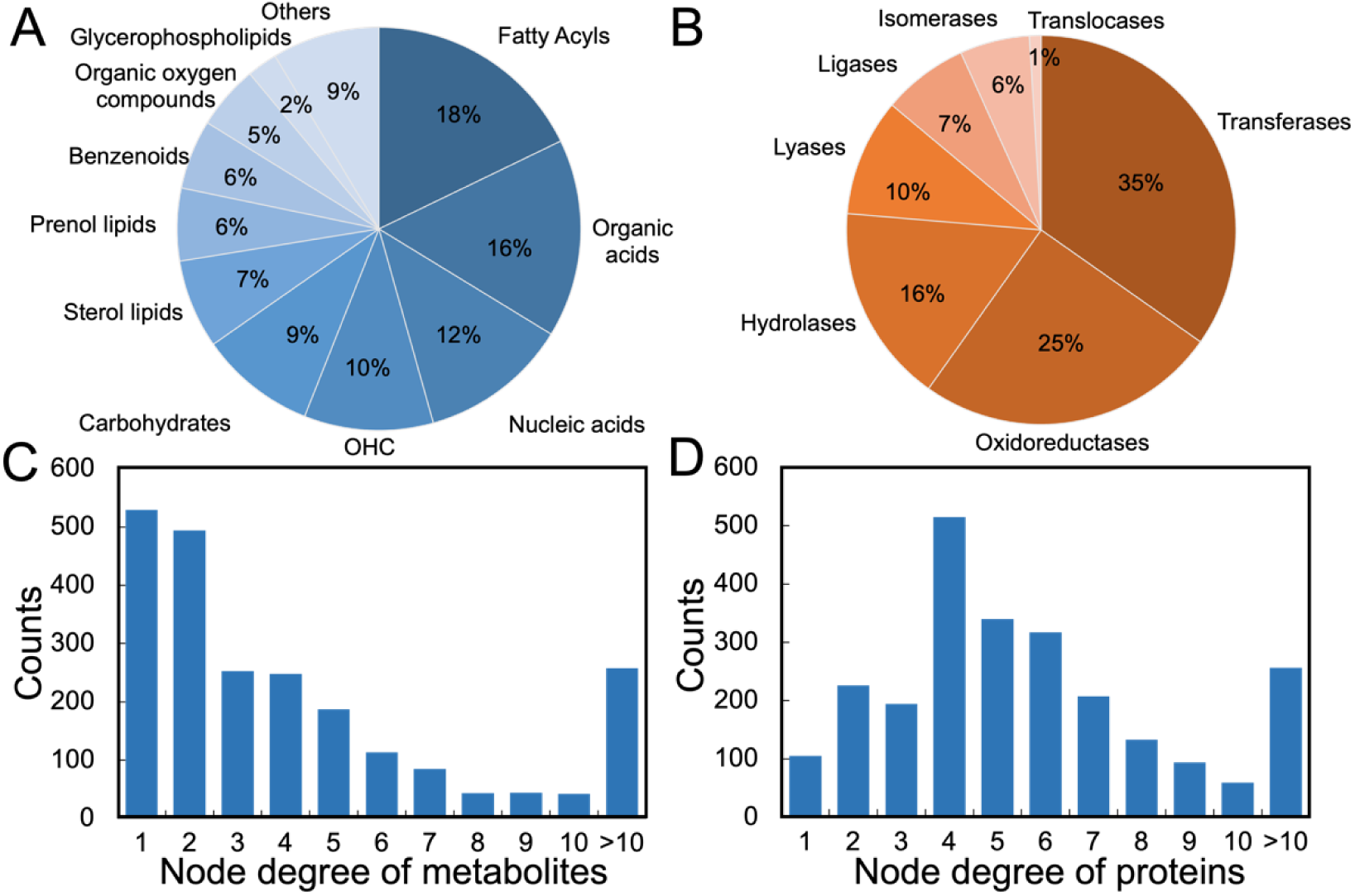
Characteristics of metabolites and proteins in the metabolite-protein interaction network. Panel A and B show the classes of metabolites and proteins in the metabolite-protein interaction network of all organisms. Panel C and D show the distributions of node degree in the metabolite-protein interaction network of all organisms.

**Figure 3.**
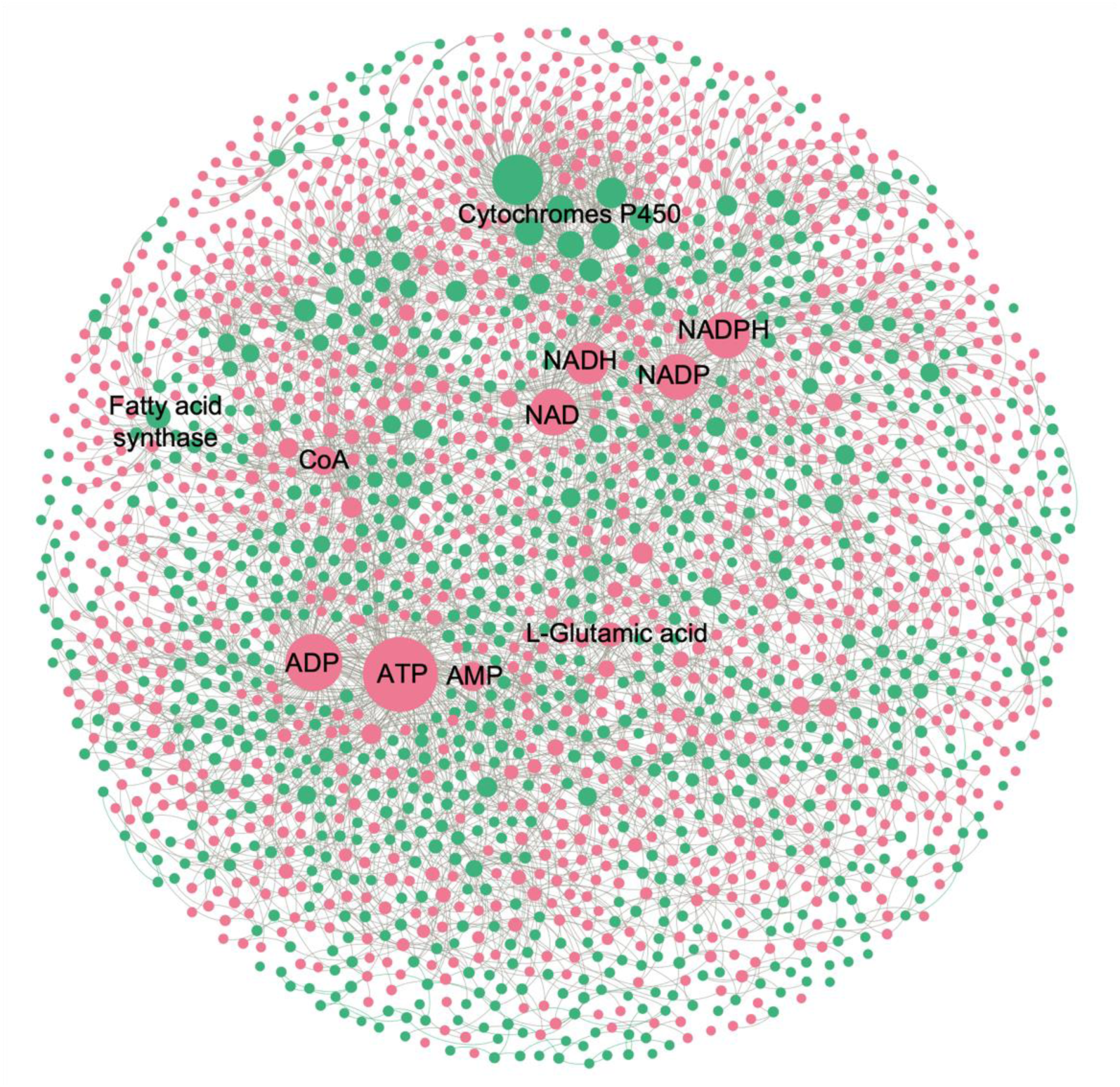
Metabolite-protein interaction network of *Homo sapiens*. The red circle denotes a metabolite and the green circle denotes a protein. The size of the circle is proportional to the node degree in the network. The top interconnected nodes are annotated on the graph.

**Table 1.** Details of the metabolite-protein interaction network across a variety of organisms.

| Organisms | Number of<br>Nodes | Number of<br>Edges | Average<br>Degree | Number of<br>Proteins | Number of<br>Metabolites |
| --- | --- | --- | --- | --- | --- |
| *Homo sapiens* | 2306 | 5786 | 5.02 | 855 | 1451 |
| *Mus musculus* | 1664 | 3826 | 4.60 | 469 | 1195 |
| *Rattus norvegicus* | 1823 | 4423 | 4.85 | 685 | 1138 |
| *Escherichia coli* | 1811 | 4288 | 4.73 | 440 | 1371 |
| *Bos taurus* | 1771 | 4240 | 4.79 | 646 | 1125 |
| *Arabidopsis thaliana* | 1286 | 2885 | 4.49 | 438 | 848 |
| *Drosophila melanogaster* | 1086 | 2568 | 4.73 | 374 | 712 |
| *Saccharomyces cerevisiae* | 1125 | 2629 | 4.67 | 308 | 817 |
| *Caenorhabditis elegans* | 1011 | 2442 | 4.83 | 322 | 689 |
| *Pseudomonas aeruginosa* | 1280 | 2935 | 4.59 | 328 | 952 |

### 3.2 Selection of feature representations of metabolites and proteins

A notable advantage of GNNs is capable to combine the node attributes with the graph topological features. The molecular structures of metabolites and proteins are critical to determine the likelihood of the enzymatic reaction. Encoding the molecular structure as the node attributes in the MPI network would enhance the performance of the VGAE model. Given that many types of numerical representations for structures of metabolites and proteins are available, we examined and selected the optimal feature representations for the MPI prediction of the VGAE model. Molecular fingerprints are a kind of fixed-length of binary vectors to represent the structures of small molecules. Importantly, they are rapid to generate and thus are selected as the representation of metabolites. Sequence-based feature extraction methods were used for proteins. To select the optimal representations of metabolites and proteins, we used different types of feature representations to train our VGAE model and compared the performances of MPI predictions as described in Methods 2.2. Here, we evaluated the models by different combinations of three molecular fingerprints and PCA transformed molecular fingerprints and two protein embedding models (SeqVec and ESM-1b transformer). In the training datasets of *Homo sapiens*, five-fold cross-validation was performed and the results of each model are summarized in Table 2. For metabolite features, it was found that PCA transformed molecular fingerprints have better performance than traditional molecular fingerprints. For proteins, the SeqVec model performed slightly better than the ESM-1b transformer. The combination of ECFP (PCA-transformed) of metabolites and SeqVec of proteins achieved the best result with an AUC score of 0.930 and an Average-Precision (AP) score of 0.938. Therefore, the ECFP (PCA-transformed) and SeqVec were selected to generate features of metabolites and proteins for model evaluation and application.

**Table 2.** Result of metabolite-protein interaction prediction of *Homo sapiens* using different representations of protein and metabolite features.

| Metabolite feature<br>(molecular fingerprint) | Protein feature | AUC score | AP score |
| --- | --- | --- | --- |
| ECFP | SeqVec | 0.921±0.002 | 0.937±0.001 |
| ECFP (PCA) | SeqVec | **0.930±0.003** | **0.938±0.002** |
| MACCS | SeqVec | 0.915±0.005 | 0.928±0.008 |
| MACCS (PCA) | SeqVec | 0.921±0.002 | 0.934±0.001 |
| Topological | SeqVec | 0.882±0.013 | 0.894±0.011 |
| Topological (PCA) | SeqVec | 0.920±0.001 | 0.938±0.004 |
| ECFP | ESM-1b | 0.914±0.005 | 0.931±0.004 |
| ECFP (PCA) | ESM-1b | 0.922±0.007 | 0.937±0.004 |
| MACCS | ESM-1b | 0.908±0.001 | 0.924±0.001 |
| MACCS (PCA) | ESM-1b | 0.917±0.001 | 0.933±0.003 |
| Topological | ESM-1b | 0.866±0.007 | 0.878±0.001 |
| Topological (PCA) | ESM-1b | 0.912±0.010 | 0.933±0.004 |

### 3.3 VGAE model performance and evaluation on organism-wise MPI networks

After selecting the optimal feature representations of metabolites and proteins, we trained the MPI-VGAE framework across all ten organisms and evaluated it on the MPI network in the testing dataset. Figure 4 shows the performance of MPI-VGAE on metabolite-protein interaction network of *Homo sapiens*. Figure 4A and 4B shows the details of training and testing result MPI-VGAE on the metabolite-protein interaction network of *Homo sapiens*. The vectorized input features and graph embeddings of metabolites and proteins were visualized by a non-linear dimensionality technique t-distributed stochastic neighbor embedding (t-SNE). t-SNE maps high-dimensional features to low-dimensional ones by reserving information during dimension reduction. Figure 4C and D shows the t-SNE visualization of feature representations and graph embeddings of metabolites and proteins of *Homo sapiens* in the VGAE model. The input features of metabolites and proteins were clearly separated as shown in Figure 4C. Since the VGAE model weighted the neighborhood node attributes in the MPI network, the graph embeddings of adjacent metabolites and proteins in the MPI network were very close (Figure 4D). Figure 4E and 4F show the comparison of ROC and PR curves between MPI-VGAE and other machine-learning models on metabolite-protein interactions of integrated MPI of *Homo sapiens*. The VGAE model embedded with structural information of metabolites and protein sequence properties obtained the highest performance (AUC: 0.915, AP: 0.931), which suggests that the node molecular features greatly enhance the performance in predicting the likelihood of MPI. Table 3 and Table S2 show all the performance of ROC and AP scores by multiple machine learning models on different organisms. The MPI-VGAE with structural information performed the best across all ten organisms. Compared with other similarity based and graph-based methods, the VGAE model with node attributes boosted the performance up to 11%, which achieved AUC scores from 0.787 to 0.924, and AP scores from 0.827 to 0.942 for the ten organisms. Due to the variance of the number of metabolic pathways among different organisms, the node degree distributions may vary across different organisms. Minor fluctuations of performance existed in different organisms due to the variation of the MPI network. For instance, the MPI network of *Arabidopsis thaliana* has the lowest average node degree and the performance of MPI prediction was the worst among all organisms. The possible reason is that the sparseness of the MPI network affects the prediction performance of MPI.

**Figure 4.**
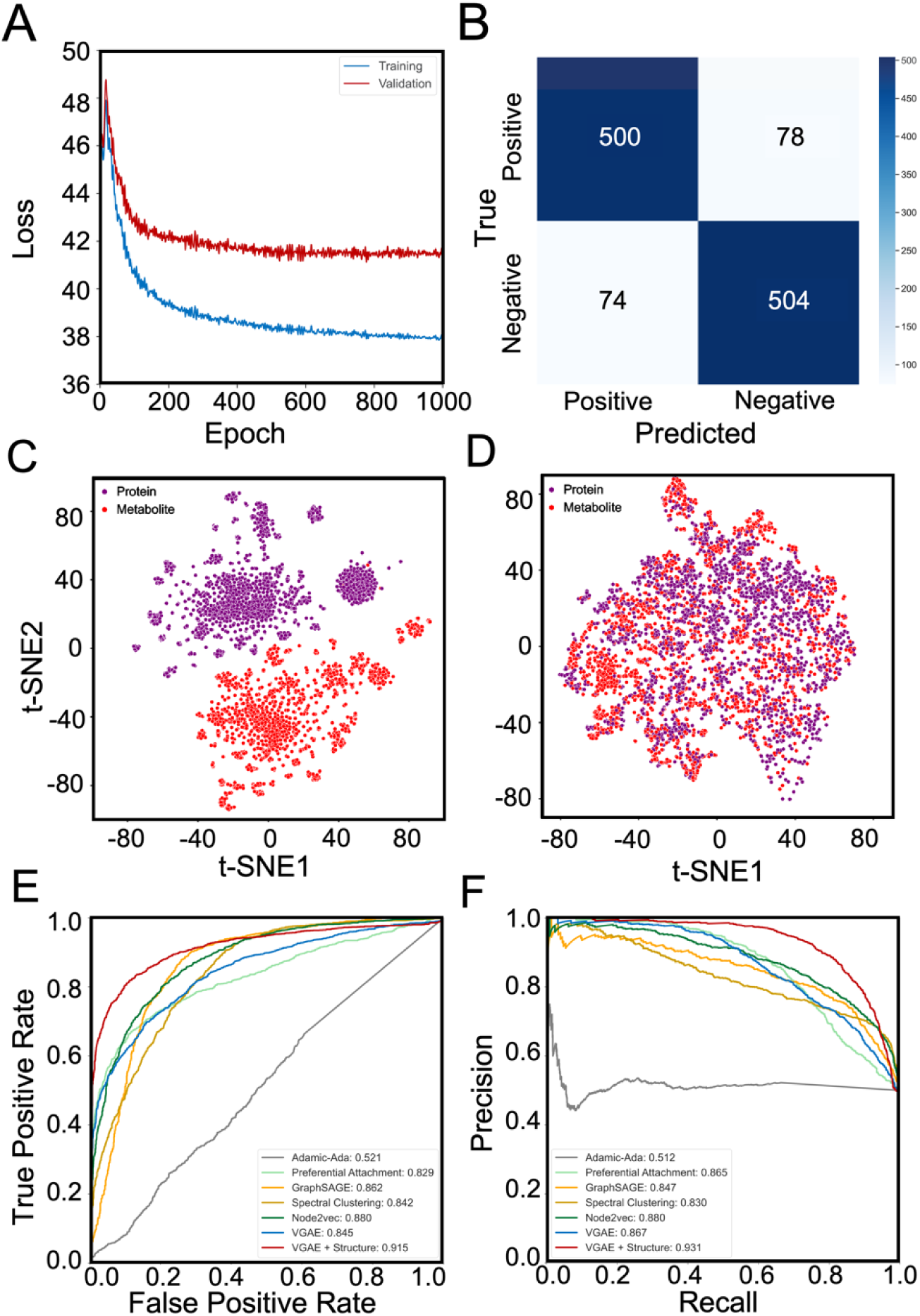
Performance of MPI-VGAE on metabolite-protein interaction network of *Homo sapiens*. Panel A shows the training and validation loss *vs.* epoch in the MPI-VGAE. Panel B shows the confusion matrix on the metabolite-protein interaction network of *Homo sapiens*. Panel C shows the t-SNE visualization of explicit feature representation by ECFP molecular fingerprints of metabolites and SeqVec features of proteins. Panel D shows the t-SNE visualization of graph embeddings of metabolites and proteins by VGAE. Panel E and F show the ROC and PR curve of MPI prediction by different machine learning models on the test dataset of the metabolite-protein interaction network of *Homo sapiens*.

**Table 3.** The AUC scores of metabolite-protein interaction prediction by different machine learning models on a variety of organisms.\

| Organism | Spectral<br>Cluster-<br>ing | Adamic-Ada | Preferential<br>Attachment | GraphSAGE | Node2vec | VGAE<br>(no struc-<br>ture) | VGAE<br>(struc-<br>ture) |
| --- | --- | --- | --- | --- | --- | --- | --- |
| *Homo sapiens* | 0.842 | 0.521 | 0.829 | 0.862 | 0.880 | 0.845 | 0.915 |
| *Mus musculus* | 0.766 | 0.472 | 0.816 | 0.867 | 0.793 | 0.814 | 0.859 |
| *Rattus norvegicus* | 0.824 | 0.573 | 0.821 | 0.821 | 0.831 | 0.788 | 0.895 |
| *Escherichia coli* | 0.741 | 0.484 | 0.753 | 0.780 | 0.728 | 0.725 | 0.787 |
| *Bos taurus* | 0.803 | 0.570 | 0.794 | 0.815 | 0.844 | 0.779 | 0.896 |
| *Arabidopsis thaliana* | 0.785 | 0.351 | 0.767 | 0.794 | 0.821 | 0.754 | 0.797 |
| *Drosophila melanogaster* | 0.748 | 0.335 | 0.793 | 0.751 | 0.796 | 0.773 | 0.851 |
| *Saccharomyces cerevisiae* | 0.763 | 0.326 | 0.778 | 0.823 | 0.825 | 0.781 | 0.835 |
| *Caenorhabditis elegans* | 0.746 | 0.330 | 0.817 | 0.844 | 0.824 | 0.788 | 0.850 |
| *Pseudomonas aeruginosa* | 0.705 | 0.416 | 0.845 | 0.853 | 0.748 | 0.810 | 0.816 |

### 3.4 Reconstruction of metabolic pathways and functional MPI networks

Next, we evaluated the ability of the MPI-VGAE models on metabolite-protein interaction prediction in complicated biological systems. The VGAE model was applied to reconstruct the metabolic pathways followed by the reconstruction of functional metabolic networks. We selected the pathways with more than five real metabolite-protein interaction pairs and generated an equal number of negative pairs randomly by considering the metabolites/proteins other than true MPI pairs in the testing dataset. There are 402 metabolic pathways covering most functional classes of metabolic pathways, such as amino acid metabolism, carbohydrate metabolism, and energy metabolism. Figure 5A-C shows the distribution of the AUC scores and AP scores of the reconstructed metabolic pathways by the VGAE model. The MPI-VGAE achieved an average AUC and AP scores of 0.928 and 0.939, respectively. For instance, there are 201 MPIs in purine metabolism. Our model could successfully predict 100% of the MPIs. For the negative MPIs randomly generated, the VGAE model also accurately predicted 84% of the MPIs as negative cases.

**Figure 5.**
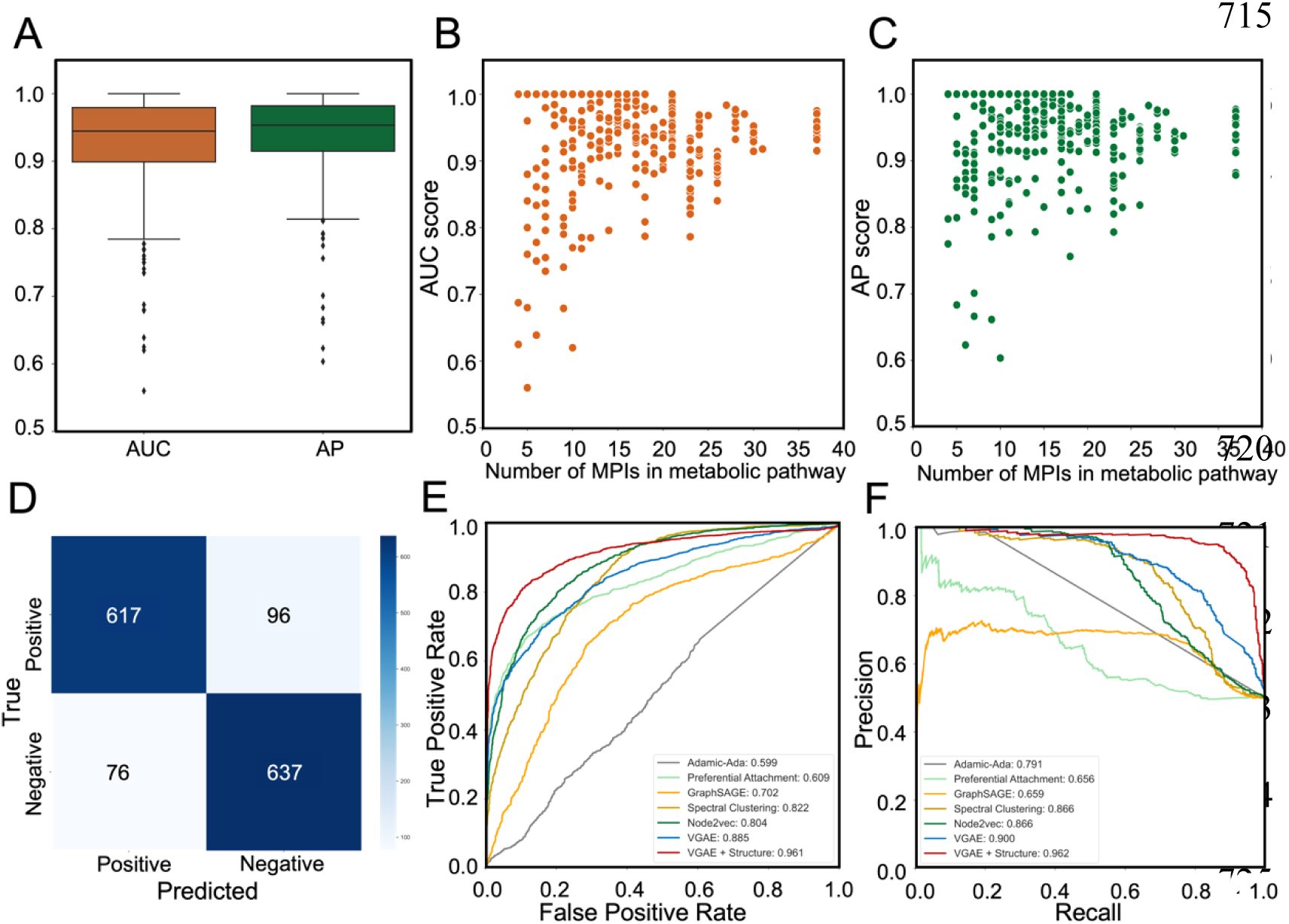
Reconstruction of metabolic pathways by the MPI-VGAE framework and applications of MPI-VGAE to KEGG metabolic reactions. Panel A shows the box plot of AUC and AP scores of metabolic pathway reconstruction by the VGAE model. To evaluate the metabolic pathway reconstruction performance of the VGAE model, 402 metabolic pathways from were included. Panel B and C show the scatter plot of AUC and AP scores *vs.* the number of MPIs in the metabolic pathway. Panel D shows the confusion matrix on the metabolic reaction network. Panel E and F show ROC and PR curve of metabolic reaction prediction by different machine learning models.

Based on the functional type of each metabolic pathway, five classes of metabolic networks were constructed, including the metabolic pathway network in biological systems, disease-associated metabolic pathway network, drug action metabolism, drug metabolism, and protein/metabolite signaling metabolic pathways. For each functional metabolic network, 80% of the positive enzymatic reactions and an equal number of negative enzymatic reactions were used for VGAE model training and optimization, and 20% of the randomized selected positive enzymatic reactions and negative enzymatic reactions were used for testing. The details of the functional metabolic networks are summarized in Table 4. The AUC score and AP score ranged from 0.883 to 0.909, and 0.899 to 0.924, showing that the performance of the VGAE model is stable across the different functional metabolic networks.

**Table 4.** Results of metabolite-protein interaction prediction based on the functional classification of MPI network.

| MPI network | Number of Nodes | Number of Edges | Average Degree | AUC score | AP score |
| --- | --- | --- | --- | --- | --- |
| Disease | 1558 | 4791 | 6.15 | 0.890 | 0.907 |
| Metabolic process | 5055 | 17506 | 6.93 | 0.883 | 0.901 |
| Drug action | 640 | 1499 | 6.68 | 0.909 | 0.924 |
| Drug metabolism | 336 | 937 | 5.58 | 0.883 | 0.899 |
| Cellular signaling | 479 | 827 | 3.45 | 0.909 | 0.927 |

### 3.5 Prediction of enzymatic reaction based on metabolic reaction network by VGAE

The metabolic reaction network is a homogenous network consisting of chemical reactions between metabolites. We evaluated the effectiveness of the VGAE model to predict the enzymatic link in the metabolic reaction network. The metabolic reaction network was constructed based on the KEGG REACTION database, which consisted of 2343 nodes with 6889 edges. We compared the performance of metabolic reaction prediction by different machine learning models and the result is shown in Figure 5D-F. By including the molecular structural information of metabolites, the VGAE model achieved an AUC score of 0.964 and an AP score of 0.962, which outperformed other similarity-based or embedding-based models such as ELP. Therefore, by embedding the structural information of nodes, the VGAE model shows excellent prediction performance in both homogeneous and heterogeneous biological networks of different organisms.

### 3.6 Application to the metabolic pathway network reconstruction in Alzheimer’s disease and colorectal cancer

Alzheimer’s disease is a progressive neurologic disorder that is the most common form of dementia. Many proteomics and metabolomics approaches have been conducted to investigate the disrupted functions of proteins and dysregulated metabolisms. We mapped the disease-associated proteins and metabolites that were induced from DisGeNET database and HMDB databases. For Alzheimer’s disease, 65 proteins and 86 metabolites were mapped to the MPI network. MPI-VGAE was applied to predict the likelihood of enzymatic reactions of all 5590 metabolite-protein interaction pairs. Eleven known existing pairs of metabolite-protein interactions such as Adenosine and Purine nucleoside phosphorylase exist in the MPI network of Alzheimer’s disease. MPI-VGAE is able to pinpoint ten out of eleven known enzymatic reactions accurately. In addition, MPI-VGAE predict eight additional enzymatic reactions with high confidence score. For instance, Cholesterol side-chain cleavage enzyme (CYP11A) has highly confident interaction with 24-Hydroxy-cholesterol. Since Cytochrome P450s (CYPs) play critical roles in cholesterol homeostasis, many CYPs were disrupted in the cholesterol metabolism and transport in AD that has been investigated by previous studies [41–44]. Due to the high structural similarity of CYPs and cholesterol derivatives, the enzymatic reaction initiated by the metabolite and protein interaction could be altered. The molecular docking simulates the interaction details between Cholesterol side-chain cleavage enzyme (CYP11A) binding with 24-Hydroxycholesterol (Figure 6B) and 27-Hydroxycholesterol (Figure S3A). We also show the molecular docking results of protein-ligand structures of protein Aldo-keto reductase family 1 member C4 (AKR1C4) binding with 27-Hydroxycholesterol (Figure 6C) and 24-Hydroxycholesterol in (Figure S3B).

**Figure 6.**
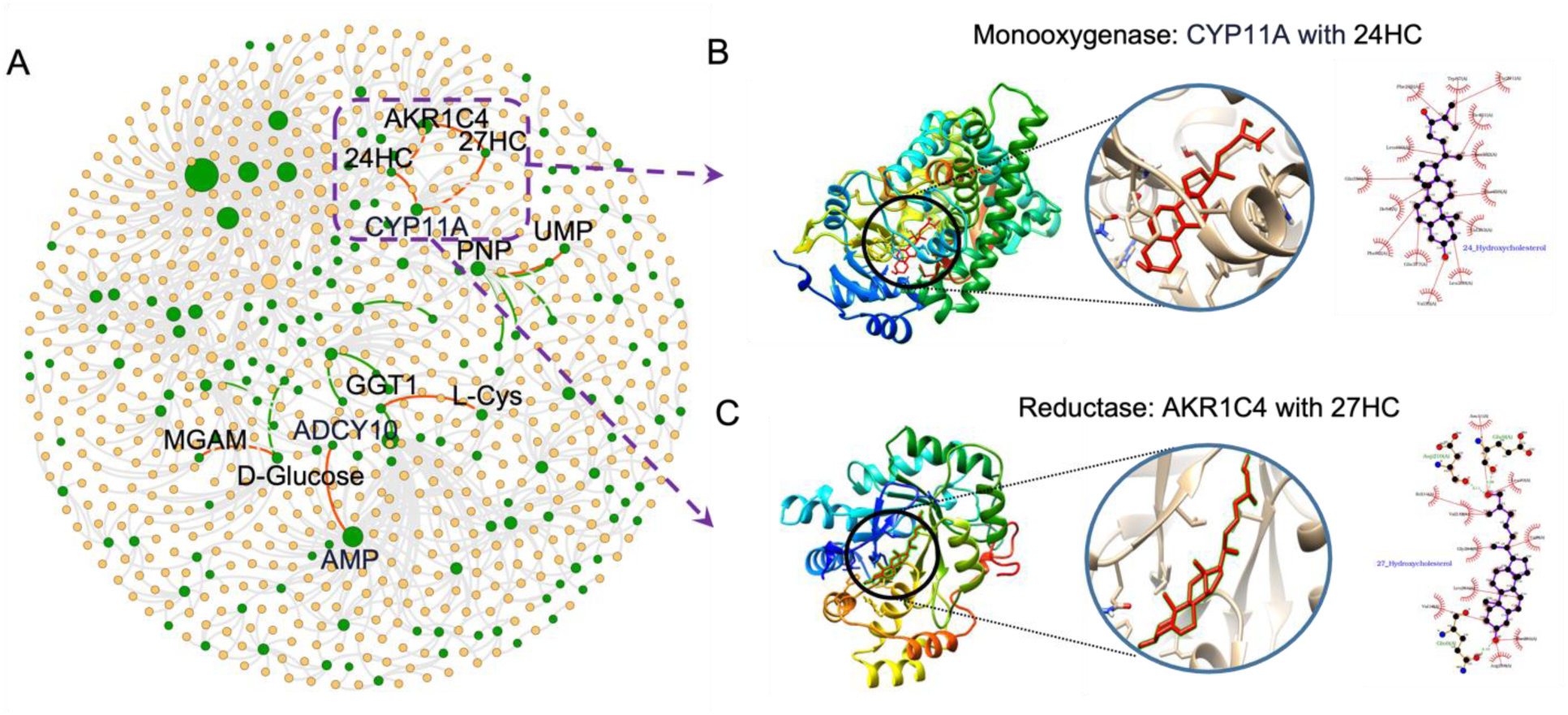
Reconstruction of MPI network of Alzheimer’s disease by the MPI-VGAE framework using 65 proteins and 86 metabolites. Panel A shows the reconstructed MPI network. The green dots denote the disrupted proteins and metabolites in Alzheimer’s disease. The orange dots denote the normal proteins and metabolites that have enzymatic reactions with the disrupted molecules. The circle size is proportional to the node degree in the MPI network. The edges denote the interaction between proteins and metabolite (gray edge: known interaction between normal proteins and metabolites, green edge: known interaction between disrupted proteins and metabolites, red edge: predicted interaction between disrupted proteins and metabolites). Panel B shows the molecular docking result between Cholesterol side-chain cleavage enzyme (CYP11A) binding with 24-Hydroxycholesterol (24HC) (binding energy:-9.6 kcal/mol). Panel C shows the molecular docking result between Aldo-keto reductase family 1 member C4 (AKR1C4) binding with 27-Hydroxycholesterol (27HC) (binding energy:-7.6 kcal/mol).

Colorectal cancer is the third-most prevalent, and second-most deadly, cancer worldwide. Integrated analysis of proteomics and metabolomics has been performed to study the proteomic and metabolic alterations in a variety of samples such as tissues and plasma. 81 proteins and 216 metabolites were gathered from DisGeNET and HMDB databases that were identified to be related to colorectal cancer. MPI-VGAE was applied to predict all the possible enzymatic reactions based on all 17496 pairs. MPI-VGAE predicts 37 out 44 known enzymatic reactions accurately. Figure S4 shows the reconstructed MPI network of enzymatic reaction prediction by MPI-VGAE for colorectal cancer. The highly confident enzymatic reactions pairs predicted by MPI-VGAE for Alzheimer’s disease and colorectal cancer are summarized in Table S3 and Table S4. In a sum, the applications to the reconstruction of metabolic reaction for Alzheimer’s disease and colorectal cancer demonstrate the efficiency and capability of MPI-VGAE to discover new disease-related enzymatic reactions and metabolic pathways.

## 4. Discussion

In this study, we have developed a graph neural network-based method to identify metabolite-protein interactions based on the MPI network. To explore the best feature representations of metabolites and proteins, we compared different combinations of feature extraction approaches based on the MPI prediction performances by the VGAE model. As shown in Table 2, the best performance was obtained by using the combination of protein features from SeqVec and metabolites features from ECFP (PCA-transformed). All the AUC-ROCs were above 0.91 except the combination with topological features of metabolites, which indicated our predictors were rather robust with different features. The implementation of the PCA method to transform the molecular fingerprints further improved the prediction performances, e.g., ESM-1b with topological features (AUC 0.788 vs 0.914). Given that PCA had learned large-scale molecular fingerprints over 78,000 metabolites, the PCA-transformed molecular fingerprints reserved both the features of metabolites and the feature variance between metabolites. In addition, the PCA-transformed molecular fingerprints were no longer binary with 0 or 1 values, which reduced the chance of gradient vanishing during the training process. Interestingly, the combination with protein features by SeqVec shows slightly better performance than features by ESM-1b transformer methods. SeqVec is more computationally efficient and widely used for the fast feature extraction of proteins. It generated the same vector length for proteins with different sequences, which might ignore the size-dependent properties of proteins.

We also tested the MPI-VGAE model to predict metabolite-protein interactions in different organisms. Our approach obtained the best performances across all organisms compared with other methods. Moreover, MPI-VGAE shows stable performance when reconstructing different functional MPI networks. As shown in Table 4, among all five categories of functional MPI networks, the AUC scores are always better than 0.88 and AP scores reach above 0.89. This indicates that the MPI-VGAE method could be able to predict the MPI network involved in the major functional classes.

A notable feature of MPI-VGAE is the capability to reconstruct the MPI network of specific disease based on a list of the disrupted metabolites and proteins. Furthermore, MPI-VGAE predict the highly likely new enzymatic reactions occurring among the metabolites and proteins, which will facilitate the understanding of disrupted metabolisms in diseases. For Alzheimer’s disease and colorectal cancer, MPI-VGAE identified a few potential new enzymatic reactions such as CYP11A and 24-Hydroxycholesterol.

Though our MPI-VGAE could achieve the best performance among all the methods, there are a couple of limitations to our study. The imbalanced distribution of positive and negative edges in the network would hamper the accurate prediction of true positive edges. For example, the high specificity but low precision would be obtained if the model predicts all MPIs as negative. To reduce the effect of the imbalanced dataset on the VGAE model, we applied the downsampling approach to generate the training and testing dataset and used AUC and AP scores for performance measures. Hopefully, with the rapidly increasing discovery of protein and metabolites interactions, the imbalanced dataset issue in the MPI network will be alleviated.

## 5. Conclusion

In this work, we present the variational graph autoencoder method to predict metabolite-protein interactions based on the MPI network. When incorporating the node attributes of metabolites and proteins, the performance of MPI-VGAE achieved the highest AUC and AP scores in different genome-scale enzymatic reaction networks. The MPI-VGAE framework also showed stable and excellent performance in reconstructing the metabolic pathways and functional MPI network. To the best of our knowledge, this is the first time that VGAE has been applied in the MPI network for efficient enzymatic reaction prediction. By applying MPI-VGAE to identify protein-metabolite reactions in Alzheimer’s disease and colorectal cancer, our method could not only find experiment-proved interactions, but also predict novel and reliable metabolite reactions which could be crucial for mechanistic investigation of disease and drug target discovery. We believe the method will greatly assist the discovery of novel disease-related enzymatic reactions and pave the way for genome-scale metabolic pathway reconstruction by graph neural network approaches.

## Competing interests

The authors declare that they have no competing interests

## Funding

This work was supported by the National Institutes of Health [R35ES028365 to G.J.P.]; and the Young Scholars Program of Shandong University [21320082164070 to C.W., 21320082064101 to Q.H.].

## Notes

https://github.com/mmetalab/mpi-vgae

### Competing Interest Statement

The authors have declared no competing interest.

